## Supplementary Material for "Correlative all-optical quantification of mass density and mechanics of sub-cellular compartments with fluorescence specificity"

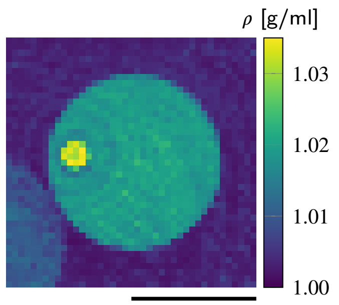


**Figure 1–figure supplement 1.** Absolute density of a cell phantom consisting of a PDMS bead inside a PAA bead. Scale bar 10 µm.

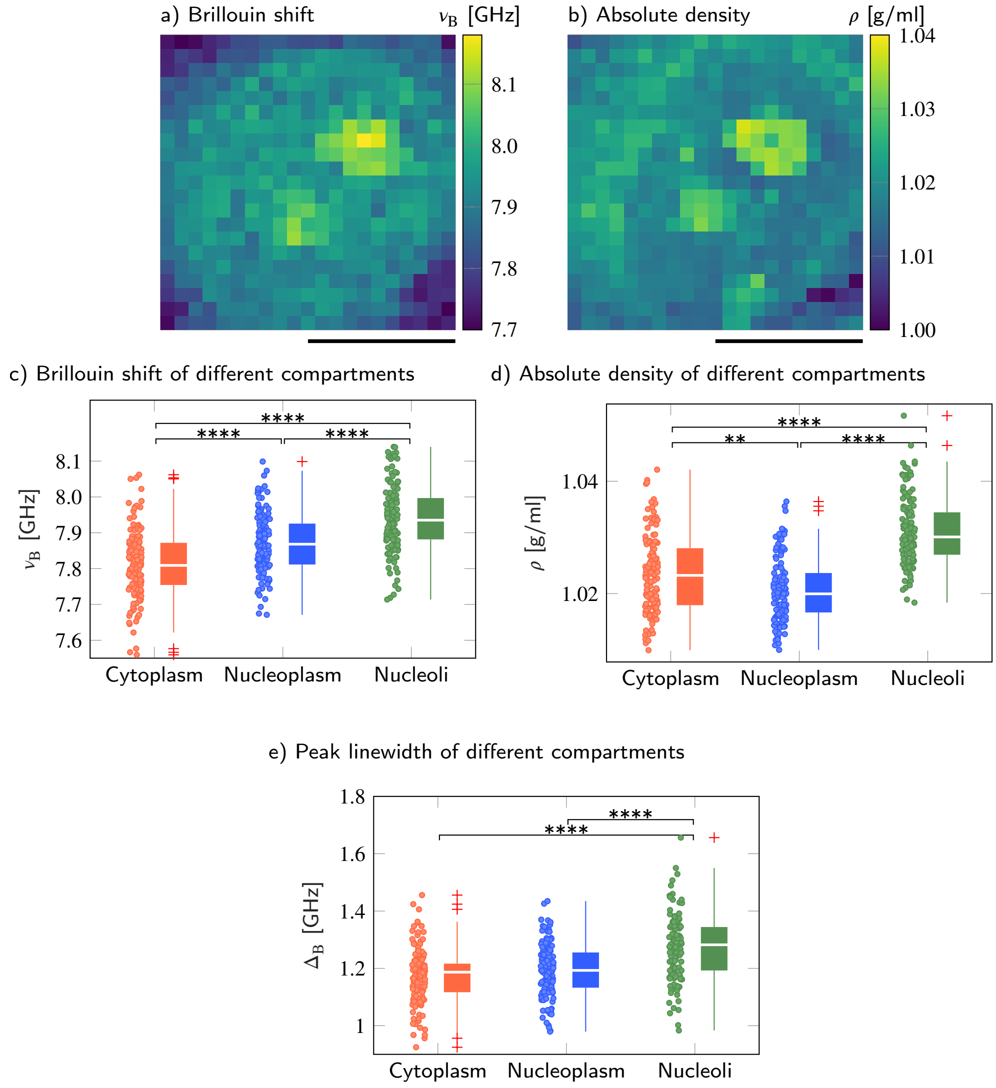


**Figure 2–figure supplement 1.** Brillouin shift and absolute density of cytoplasm, nucleoplasm and nucleoli in HeLa cells. Representative maps of the (**a**) Brillouin shift and (**b**) absolute density of a HeLa cell. Quantitative analysis of (**c**) the Brillouin shift $\nu_B$, (**d**) the absolute density $\rho$ and (**e**) peak linewidth $\Delta_B$. **$p < 0.01$; ****$p < 0.0001$. Scale bars $10\,\mu$m.

**Supplementary File 1.** Average values and standard errors of the mean of the RI $n$, Brillouin shift $\nu_B$, absolute density $\rho$, longitudinal modulus $M'$ and linewidth $\Delta_B$ for the cytoplasm (cyto), nucleoplasm (np) and nucleoli (nl) of 139 wild-type HeLa cells.

| compartment | RI $n$ | Brillouin shift $\nu_B$ [GHz] | absolute density $\rho$ [g/ml] | longitudinal modulus $M'$ [GPa] | linewidth $\Delta_B$ [GHz] |
|---|---|---|---|---|---|
| cytoplasm | $1.3545 \pm 0.0004$ | $7.811 \pm 0.008$ | $1.0234 \pm 0.0006$ | $2.410 \pm 0.005$ | $1.175 \pm 0.008$ |
| nucleoplasm | $1.3522 \pm 0.0004$ | $7.872 \pm 0.007$ | $1.0207 \pm 0.0005$ | $2.448 \pm 0.005$ | $1.193 \pm 0.008$ |
| nucleoli | $1.3618 \pm 0.0004$ | $7.938 \pm 0.008$ | $1.0310 \pm 0.0005$ | $2.487 \pm 0.005$ | $1.271 \pm 0.010$ |

**Supplementary File 2.** Kruskal-Walis $p$-values when comparing the RI $n$, Brillouin shifts $\nu_B$, mass densities $\rho$, longitudinal moduli $M'$ and linewidths $\Delta_B$ of the cytoplasm (cyto), nucleoplasm (np) and nucleoli (nl) of 139 wild-type HeLa cells, respectively.

| comparing | RI<br>$p_{n_{c_1},n_{c_2}}$ | Brillouin shift<br>$p_{\nu_{B,c_1},\nu_{B,c_2}}$ | absolute density<br>$p_{\rho_{c_1},\rho_{c_2}}$ | longitudinal modulus<br>$p_{M'_{c_1},M'_{c_2}}$ | linewidth<br>$p_{\Delta_{B,c_1},\Delta_{B,c_2}}$ |
|---|---|---|---|---|---|
| $c_1$:cyto to $c_2$:np | $9 \times 10^{-4}$ | $2 \times 10^{-6}$ | $3 \times 10^{-3}$ | $7 \times 10^{-7}$ | 0.139 |
| $c_1$:cyto to $c_2$:nl | $1 \times 10^{-21}$ | $1 \times 10^{-23}$ | $7 \times 10^{-17}$ | $4 \times 10^{-23}$ | $8 \times 10^{-14}$ |
| $c_1$:np to $c_2$:nl | $2 \times 10^{-37}$ | $2 \times 10^{-7}$ | $3 \times 10^{-29}$ | $9 \times 10^{-7}$ | $2 \times 10^{-8}$ |

**Supplementary File 3.** Average values and standard errors of the mean of the RI $n$, Brillouin shift $\nu_B$, absolute density $\rho$ and longitudinal modulus $M'$ for the cytoplasm and polyQ aggregates of 22 wild-type HeLa cells.

| compartment | RI<br>$n$ | Brillouin shift<br>$\nu_B$ [GHz] | absolute density<br>$\rho$ [g/ml] | longitudinal modulus<br>$M'$ [GPa] |
|---|---|---|---|---|
| cytoplasm | $1.3506 \pm 0.0013$ | $7.861 \pm 0.014$ | $1.020 \pm 0.002$ | $2.442 \pm 0.009$ |
| polyQ aggregate | $1.3856 \pm 0.0018$ | $8.789 \pm 0.040$ | $1.061 \pm 0.003$ | $3.051 \pm 0.029$ |

**Supplementary File 4.** Average values and standard errors of the mean of the RI $n$ and longitudinal modulus $M'$ for different conditions and compartments of P525L HeLa cells.

| condition | compartment | RI<br>$n$ | longitudinal modulus<br>$M'$ [GPa] |
|---|---|---|---|
| control | cytoplasm | $1.3481 \pm 0.0010$ | $2.361 \pm 0.009$ |
| | nucleoplasm | $1.3509 \pm 0.0008$ | $2.409 \pm 0.006$ |
| arsenite | peri-SG cytoplasm | $1.3456 \pm 0.0007$ | $2.329 \pm 0.009$ |
| | nucleoplasm | $1.3476 \pm 0.0006$ | $2.394 \pm 0.009$ |
| | SGs | $1.3443 \pm 0.0006$ | $2.345 \pm 0.009$ |
| arsenite, fixed | peri-SG cytoplasm | $1.3443 \pm 0.0005$ | $2.331 \pm 0.006$ |
| | nucleoplasm | $1.3485 \pm 0.0005$ | $2.416 \pm 0.006$ |
| | SGs | $1.3454 \pm 0.0004$ | $2.357 \pm 0.006$ |

### Localized measurements do not affect cell viability

In order to analyze the influence of the laser exposure due to Brillouin microscopy on the viability of P525L HeLa cells which express GFP-tagged FUS, we compared different acquisition schemes and laser wavelengths. With the FOB setup using a laser with a wavelength of 532 nm we performed acquisitions scanning the whole cell (Figure 4–figure supplement 1 first column) and a small region of 5 µm by 5 µm in the cytoplasm (Figure 4–figure supplement 1 second column). With a stand-alone Brillouin microscope which uses a laser with a longer wavelength of 780 nm (described in ***Schlüßler et al.*** (***2018***)) we performed a comparative measurement also scanning the complete cell (Figure 5–figure supplement 1 third column). We acquired fluorescence and brightfield images before and after Brillouin acquisition. Furthermore, after Brillouin acquisition, the cells were incubated for 10 min at room temperature with serum-free Dulbecco's modified Eagle's medium (DMEM) (31966-021, Thermo Fisher) supplemented with 8 µg/ml fluorescein diacetate (FDA) and 20 µg/ml propidium iodide (PI) (Sigma-Aldrich) to check the cell viability.

We find that after scanning the whole cell with the FOB setup, the fluorescence intensity in the GFP channel of the HeLa P525L cell measured is reduced due to photo bleaching (Figure 4–figure supplement 1j) and the cell starts to bleb (Figure 4–figure supplement 1m). Furthermore, the cell probed shows no enzymatic activity anymore and PI can enter the nucleus and nucleoi. When only probing a small region around a stress granule, the cell selected shows no signs of blebbing (Figure 4–figure supplement 1n) after the measurement using the FOB setup, enzymatic activity is present and PI does not enter the nucleus (Figure 4–figure supplement 1q). When using a longer wavelength of 780 nm the cell tested appears unaffected of the Brillouin measurement, shows a similar fluorescence intensity compared to before the Brillouin acquisition (Figure 4–figure supplement 1l), no blebbing (Figure 4–figure supplement 1o) and enzymatic activity (Figure 4–figure supplement 1r). Furthermore, wild-type HeLa cells were substantially less susceptible to laser illumination than P525L HeLa cells expressing GFP-tagged FUS and showed no blebbing even after a whole cell acquisition.

In summary, this shows that the cells presented in the manuscript are viable after Brillouin acquisition and the FOB setup can be considered an *in vivo* technique. However, a longer wavelength of 780 nm would have positive effects on the cell viability and, hence, should be used for future realizations of the technique.

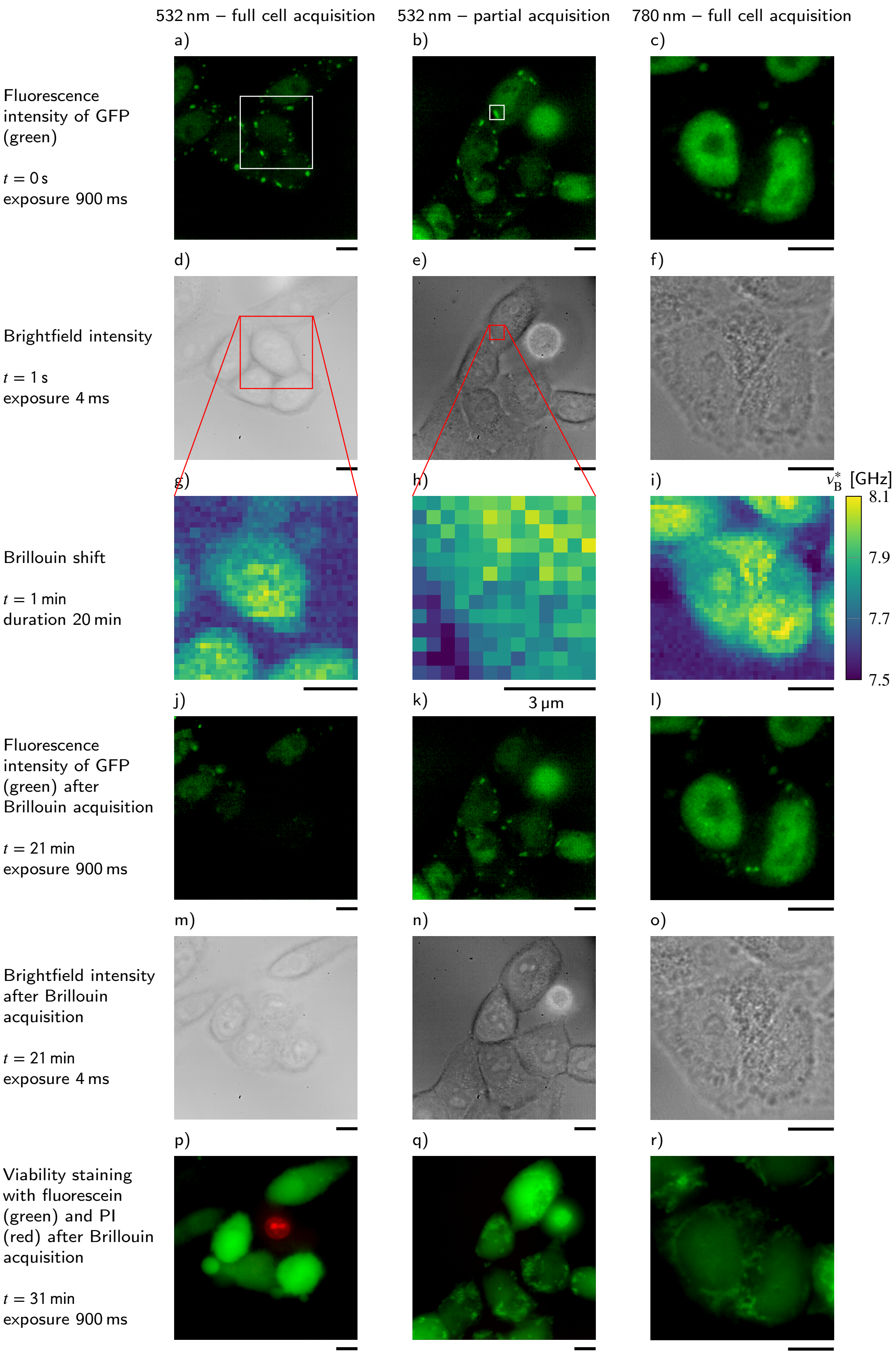


**Figure 4–figure supplement 1.** Comparison of the influence of different acquisition schemes and laser wavelengths on the viability of P525L HeLa cells which express GFP-tagged FUS. For better comparability the Brillouin shift $\nu_B^*$ measured with the $780\,\mathrm{nm}$ setup is normalized to a wavelength of $532\,\mathrm{nm}$. Scale bars $10\,\mu\mathrm{m}$.

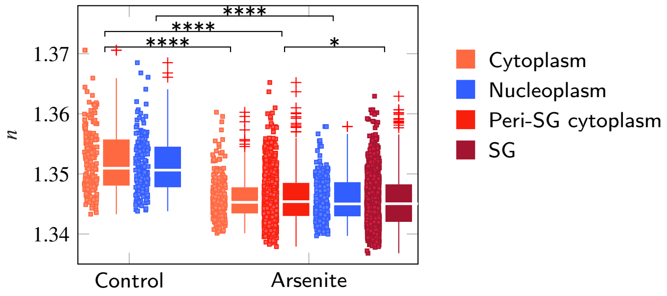


**Figure 4–figure supplement 2.** Evaluation of the RI of P525L FUS HeLa cells taking into account the complete cell. FUS-GFP-labelled stress granules induced by oxidative stress show a similar RI as the peripheral cytoplasm. *$p < 0.05$; ****$p < 0.0001$.
